# Faces under continuous flash suppression capture attention faster than objects, but without a face evoked steady-state visual potential: Is curvilinearity responsible for the behavioral effect?

**DOI:** 10.1101/778373

**Authors:** Andrew D. Engell, Henry Quillian

## Abstract

Face perception is a vital part of human social interactions. The social value of faces makes their efficient detection evolutionarily advantageous. It has been suggested that this might occur nonconsciously, but experimental results are equivocal thus far. Here, we probe nonconscious face perception using a novel combination of binocular rivalry with continuous flash suppression, and steady-state visually evoked potentials. In the first two experiments, participants viewed either non-face objects, neutral faces (Study 1), or fearful faces (Study 2). Consistent with the hypothesis that faces are processed nonconsciously, we found that faces broke through suppression faster than objects. We did not, however, observe a concomitant face-selective SSVEP. Study 3 was run to reconcile this paradox. We hypothesized that the faster breakthrough time was due to a mid-level visual feature, curvilinearity, rather than high-level category membership, which would explain the behavioral difference without neural evidence of face-selective processing. We tested this hypothesis by presenting participants with four different groups of stimuli outside of conscious awareness: rectilinear objects (e.g., chessboard), curvilinear objects (e.g., dartboard), faces, and objects that were not dominantly curvilinear or rectilinear. We found that faces *and* curvilinear objects broke through suppression faster than objects and rectilinear objects. Moreover, there was no difference between faces and curvilinear objects. These results support our hypothesis that the observed behavioral advantage for faces is due to their curvilinearity, rather than category membership.

**Highlights:**

- Faces presented outside of awareness do not evoke a steady-state visually evoked potential.
- This is true for both neutral and fearful faces.
- However, faces do breakthrough interocular suppression faster than objects.
- Curvilinear objects breakthrough interocular suppression faster than rectilinear objects.
- The breakthrough time advantage for faces over objects is due to their curvilinearity.

## 1. Introduction

Faces are considered to be a special category of visual stimuli. In this view, the social and behavioral importance of these ubiquitous stimuli created an evolutionary pressure that resulted in sensory-cognitive processes and neural machinery specialized for face perception. But how special is special? The limited processing of the visual system necessarily means that some stimuli, particularly those outside of attentional focus and awareness, will only be processed superficially. Are faces equally vulnerable to this superficial treatment by the visual system, or does their evolutionary importance result in more complete processing even when presented outside of awareness? The latter is an intuitively appealing notion, but empirical support has been equivocal (Axelrod, Bar, & Rees, 2015).

One approach to investigating nonconscious processing relies on the interocular suppression that occurs when each eye views a different image (binocular rivalry). Visual awareness will alternate between the stimuli, such that the initially suppressed image will reach awareness and vice versa. Continuous flash suppression (CFS) is a type of binocular rivalry paradigm that extends the potential duration of the suppression from seconds to minutes (Tsuchiya & Koch, 2005). Though CFS dramatically increases the duration of suppression, the suppressed images will eventually break through into awareness. Breakthrough of continuous flash suppression (b-CFS) paradigms leverage this by inferring differences in nonconscious processing if breakthrough times systematically vary across conditions (Jiang, Costello, & He, 2007).

This approach can be particularly powerful when paired with magneto-/electro-encephalography (M/EEG), which can potentially yield an objective and temporally high-resolution electrophysiological marker of face-selective processing. However, this approach has yet to yield conclusive evidence, one way or the other, of selective nonconscious face processing. Several studies have reported an increased face-related response during nonconscious detection or discrimination of neutral faces, emotive faces, or inverted faces (Jiang & He, 2006; Jiang et al., 2009; Sterzer, Jalkanen, & Rees, 2009; Suzuki & Noguchi, 2013; Baroni et al., 2017; Heering, Beauny, Vuillaume, Salvesen, & Cleeremans, 2019), but several others have found no such evidence (Reiss & Hoffman, 2007; Harris, Wu, & Woldorff, 2011; Navajas, Ahmadi, & Quian Quiroga, 2013; Shafto & Pitts, 2015; Kume et al., 2016). The inconsistent findings across studies can potentially be attributed to one or more methodological issues. These include: 1) inconsistent power to detect face-selective EEG signals during nonconscious processing, 2) different blinding methods, and 3) variation in how each study operationalizes “awareness”.

In the current series of EEG and behavioral experiments, we investigate nonconscious face processing using a novel combination of methods in an effort to address the potential limitations of prior work. Specifically, we record steady-state visually evoked potentials (SSVEP) while presenting faces and objects in a binocular rivalry with CFS paradigm. Relative to other EEG analysis techniques (e.g., ERP), SSVEP has high SNR (Norcia, Appelbaum, Ales, Cottereau, & Rossion, 2015). SSVEP relies on the periodicity of the entire dataset and should, therefore, be less susceptible to individual differences in detection criterion. For example, a conservative detection criterion that results in a short duration in which consciously perceived faces are unreported should not result in a Type I error (i.e., an SSVEP that appears to support nonconscious processing due to the contribution of a brief period of conscious perception).

Here, we test the hypothesis that face processing occurs without the benefit of conscious awareness. In the first two studies, we used a novel combination of CFS and SSVEP to look for a face-selective response when faces were presented outside of awareness. Our predictions were two-fold: that faces would breakthrough CFS faster than non-faces, and that we would observe a face-selective SSVEP. Our results were inconsistent in that we observed the former, but not the latter. To reconcile this paradox, we report a third study in which we investigated whether a midlevel feature of faces – curvilinearity – was responsible for the faster breakthrough time, rather than high-level category membership, and thus the lack of face-selective neural signature.

## 2. Methods

### 2.1. Participants

Participants were recruited from the Kenyon College campus and surrounding community and compensated for their participation. All participants had normal or corrected-to-normal vision. All participants gave written and informed consent. The Kenyon College Institution Research Board approved this protocol.

#### 2.1.1. Participants: Study 1

Data were collected from 30 participants. Five participants were excluded from analysis. Two were excluded because they had previously seen a pilot version of the paradigm. One was excluded for failing to follow instructions, one for poor data quality, and another due to equipment failure. The remaining 25 participants included 11 men and 14 women with a median age of 21.

#### 2.1.2. Participants: Study 2

Data were collected from 36 participants. Eleven participants were excluded from analysis. Four were excluded because they had previous experience with the paradigm. Three were excluded due to equipment failure, one for poor data quality, two for failing to follow instructions, and one because of a metal plate in their skull. Finally, one was excluded because they experienced immediate breakthrough in all of the CFS conditions, indicating that the binocular rivalry was completely ineffective in achieving interocular suppression. The remaining 24 participants included 7 men and 17 women with a median age of 21.

#### 2.1.3. Participants: Study 3

Data were collected from 41 participants. Six participants were excluded for disregarding or misunderstanding instructions. The remaining 35 participants included 12 men and 25 women with a median age of 21.

### 2.2. Stimuli

We converted all images to greyscale, 200 × 200 jpegs with a resolution of 72 pixels per inch. We then matched the mean luminance across images (M = 135, SD = 45) with the SHINE MATLAB toolbox (Willenbockel et al., 2010) and added a cyan filter (00FFFF) using Photoshop CS6 (see Figure 1).

**Figure 1.**
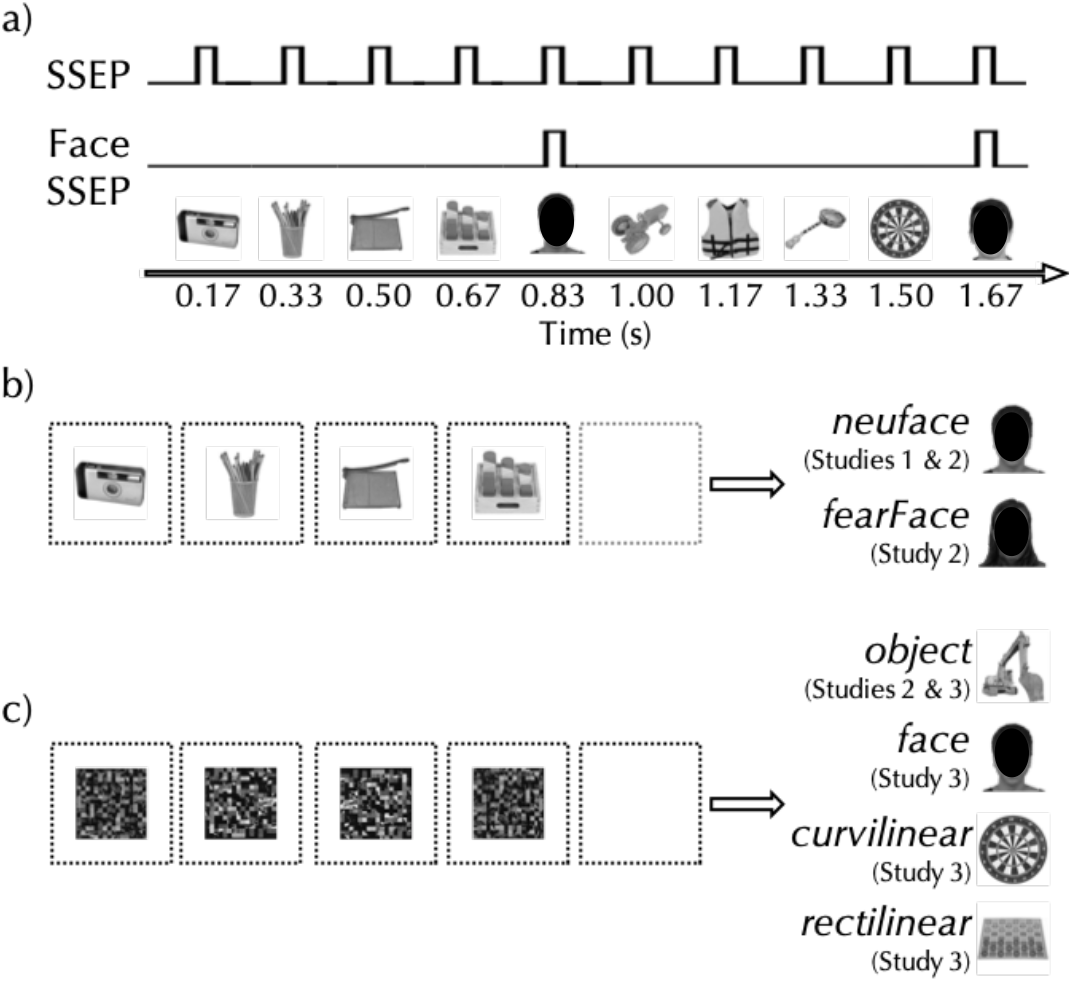
Stimuli and Paradigm: (*Face images are obscured for this preprint*). A schematic example of the paradigm and hypothetical neural response (a). Stimuli are presented periodically every 167 ms. Most of these images are drawn from one category (e.g., objects), but every 5^th^ stimulus is drawn for a different category (e.g., faces). Thus, image presentation (regardless of category) is fixed at 6 Hz, whereas the ‘oddball’ stimuli (e.g., faces) are presented at one-fifth that rate, 1.2 Hz. Across the three studies, there were a total of six unique conditions. In Study 1, the *neuFace* condition (a) displayed common objects as the frequent stimuli and neutral faces as the oddballs. Study 2 included this same condition but added *fearFace* (b), in which faces displaying a fearful expression were the oddballs. Study 2 also included the *object* condition (c), in which scrambled images were displayed as the frequent stimuli and images of objects were the oddballs. Study 3 included four conditions that all used scrambled images as frequent stimuli (c). Each of the four conditions used images from different categories as the oddballs: objects in the *object* condition (as used in Study 2), neutral faces in the *face* condition, objects dominated by curvilinear edges in the *curvilinear* condition, and objects dominated by rectilinear edges in the *rectilinear* condition.

Continuous flash suppression stimuli consisted of a 20 × 20 matrix of 20 px^2^ cells. Every 166.67 ms the cells cell colors (red or white) would be re-randomized This code was adapted from the code available at https://perso.univ-lyon2.fr/~brogniar/notes/psychopy-continuous-flash/#head.flash_init.exp. A video demonstration of the paradigm is available at https://osf.io/ysx8e/.

### 2.3. Experimental Procedure

Stimulus presentation was controlled by PsychoPy (Peirce et al., 2019) and images were displayed on a 27” LCD display with a resolution of 1920 × 1080 and a refresh rate of 0 Hz. Participants were seated ~70 cm from the display; the exact distance varied to accommodate participant comfort. Interocular suppression was achieved using red-cyan anaglyph glasses, which the participants wore throughout the experiment.

All trials began with an instruction screen that directed participants to press a button the moment they became aware of any images other than the CFS. The participant began the experiment with a button press, at which point a red fixation cross, centered within a black 500 × 500 px frame appeared at the center of the display. The frame and fixation cross remained onscreen throughout the presentation. The CFS began after three seconds, and the presentation of stimuli began two seconds after that. Images were presented at 400 × 400 px. Opacity was reduced from 50% (which would be the maximum for double exposure with the continuous flash) to 20% to facilitate suppression. Each image remained on screen for 166.67 ms (10 frames at 16.67 ms per frame refresh), with an oddball stimulus presented as every fifth image. Thus, image presentation was at 6 Hz, whereas oddball presentation was at 1.2 Hz (Figure 1). This image presentation timing was modeled after several recent reports (see Norcia et al., 2015). Moreover, continuous flash has been found most effective at achieving suppression at frequencies less than 10 Hz (Han, Lunghi, & Alais, 2016; Han et al., 2016; Zhan, Engelen, & de Gelder, 2018), particularly at or around 6 Hz (Zhu, Drewes, & Melcher, 2016; Zhan et al., 2018). Presentation would terminate after 50 cycles of four non-oddball and one oddball image or upon the participant indicating awareness of the suppressed images. The presentation order of the conditions was randomized across participants, but the three runs per condition were always presented sequentially. In Studies 1 and 2, there were complimentary conditions during which there was no CFS (*noCFS*) and therefore all images were consciously perceived. These *noCFS* condition runs were always presented at the end of the experimental session after the participant had completed all of the *CFS* conditions.

Note, the stimulus presentation rate and the flash rate were both 6 Hz. For studies 1 and 2, this means that any SSVEP to frequent image presentation was confounded with any SSVEP response to the CFS. We accepted this limitation because prior work (Han & Alais, 2018), and our pilot study observations, showed that suppression is most effective when the CFS frequency is matched to the stimulus presentation frequency. Critically, SSVEP to the oddball stimuli of interests is independent of any response to the CFS. The sole exception would be common harmonics of the oddball and CFS SSVEPs (e.g., the fourth harmonic of the oddball frequency is the same as the CFS fundamental frequency), but these were not included in analysis.

#### 2.3.1. Experimental Procedure: Study 1

Study 1 was composed of two conditions, *neuFace*, in which objects were displayed as the frequent stimuli and neutral faces as the oddball stimuli, presented either with or without continuous flash suppression (e.g., “*neuFace noCFS*”). Object and face images came from the set made available by Brady, Konkle, Alvarez, and Oliva (2008) and the “MR2 Face Database” (Strohminger et al., 2016), respectively. We selected 200 object images, excluding those that suggested animacy (e.g., dolls, toy animals) or any with a face-like appearance. We randomly selected 50 of the 74 faces available in the MR2 Face database.

#### 2.3.2. Experimental Procedure: Study 2

Study 2 was composed of six conditions: *neuFace, fearFace*, and *object*, each presented either with or without continuous flash suppression (e.g., “*neuFace noCFS*”). The *neuFace* condition was identical to the *neuFace* condition of Study 1. The *fearFace* condition displayed objects as the frequent stimuli and fearful faces as the oddball stimuli. Fearful faces oriented directly forward were taken from The Averaged Karolinska Directed Emotional Faces (Lundqvist & Litton, 1998) stimulus set. In the *object* condition, grid scrambled objects were displayed as the frequent stimuli and objects as the oddball stimuli. Images were scrambled in MATLAB by dividing the image into a 20 × 20 matrix and then randomly shuffling the location of each cell in the matrix.

#### 2.3.3. Experimental Procedure: Study 3

Study 3 was composed of four conditions: *face, object, curvilinear*, and *rectilinear.* In all conditions, scrambled objects were displayed as the frequent stimuli. In the *fa*ce condition, oddball images were seven neutral face images from the MR2 set described above. In the *object* condition, oddball images were seven common objects (e.g., backhoe) that were selected for not being dominantly curvilinear or rectilinear. In the *curvilinear* condition, oddball images were seven common curvilinear objects (e.g., dartboard). In the *rectilinear* condition, oddball images were seven common rectilinear objects (e.g., chessboard).

### 2.4. EEG Acquisition and Preprocessing

Continuous biopotential signals were recorded using the ActiveTwo BioSemi amplifier system (BioSemi, Amsterdam, The Netherlands). EEG was acquired from 64 scalp electrodes arranged in the 10/20 system. Two external electrodes were placed on the mastoids to be used as an offline reference. Two external electrodes were placed approximately 1 cm lateral and 1 cm inferior to the outer canthus of the left eye to record the horizontal and vertical electrooculogram (EOG), respectively.

All signals were digitized and recorded on an Apple Mac Mini running ActiView software (BioSemi) at a sampling rate of 2048 Hz. Off-line preprocessing and analysis were conducted with the EEGLAB (Swartz Center for Computational Neuroscience, La Jolla, CA, USA), and LETSWAVE6 (https://www.letswave.org/) MATLAB toolboxes, respectively.

Data were imported into EEGLAB, downsampled to 256 Hz, and bandpass filtered with a 4th order Butterworth filter with cutoffs of .01 – 100 Hz. Data were then cropped to only include the 41.67 s of stimulation plus an additional 1 s window before and after. For each run, the PREP pipeline (Bigdely-Shamlo, Mullen, Kothe, Su, & Robbins, 2015) was used to identify and interpolate bad channels and establish a “true – average reference. Runs in which more than ten channels required interpolation were excluded from subsequent analysis. In Study 1, 24% and 20% of runs were excluded from the CFS and noCFS conditions, respectively. In Study 2, the range of excluded runs across all six conditions was 10.64 – 18.31%.

### 2.5. Analysis

#### 2.5.1. Analysis: Behavior

The breakthrough time during continuous flash suppression was compared to the maximum run duration by subtracting the former from the latter. Therefore, a larger value indicates a faster breakthrough of interocular suppression. In Study 1, we evaluated whether the breakthrough time was greater than zero using a one-sided one-sample *t*-test. In Studies 2 and 3, we evaluated whether the breakthrough time varied across conditions with one-way repeated-measures ANOVAs and Bonferroni corrected pairwise comparisons. The Greenhouse–Geisser correction was used to correct for any violations of sphericity. ANOVA results were explicated with one-way paired-samples *t*-tests.

#### 2.5.2. Analysis: EEG

The preprocessed data were imported into LETSWAVE6 (https://www.letswave.org/) and segmented into epochs that included twelve full cycles (10 s), starting with the third image of the second cycle and ending with the second image of the twelfth cycle. For complete 50-cycle runs (e.g., those without flash suppression), this resulted in four 12-cycle epochs per run. For runs that were terminated early due to CFS breakthrough, the maximum number of non-overlapping 12-cycle epochs were extracted and the remainder discarded. The decision to discard remainder cycles was motivated by the need for sufficient frequency resolution. A 12-cycle run yields a frequency resolution of 0.1 Hz (f resolution = 1/duration = 1/10 = .01 Hz).

The 12-cycle epochs were averaged for each participant and condition. In an effort to match the SNR of the *noCFS* and *CFS* conditions, the number of epochs included in each participant’s condition averages was determined by the maximum number of available epochs in the *CFS* condition.

After discarding participants with fewer than one full cycle and runs with an excessive number of noisy channels, the following sample sizes were available for SSVEP analysis. In Study 1: *neuFace* (N=19). In Study 2: *neuFace* (N=20), *fearFace* (N=19), *object* (N=20). Note, these Study 2 samples represent subsets of the same 22 participants, with 17 participants in common across all conditions.

A fast-Fourier transform (FFT) was applied to the average time-series of 12 cycles for each participant and condition. The results were then baseline corrected by subtracting the surrounding 16 bins (8 bins on each side) excluding the local maximum and minimum. We chose 8 bins on each side to avoid contribution from neighboring harmonics, which occurred at multiples of 1.2 Hz, or 12 bins with our frequency resolution of .1 Hz. To facilitate visualization, each bin was z-normalized relative to the same range of bins described above.

We visually inspected the scalp distribution of power at the face evoked frequency for the *neuFace-noCFS* conditions and found the largest response at electrodes over the right occipitotemporal scalp: P8, PO8, and P10 (see Figures 3 and 4). This is consistent with several prior SSVEP studies of face perception (Ales, Farzin, Rossion, & Norcia, 2012; Boremanse, Norcia, & Rossion, 2013; Liu-Shuang, Norcia, & Rossion, 2014) and so these electrodes were selected as the region of interest for subsequent analysis. The statistical tests described below were run on the average of the first and second harmonics averaged across all three sites.

We used Bayesian one-sample *t*-tests (Jeffreys, 1961) as implemented in JASP 0.10.2 (JASP Team, 2019) to test whether the SSVEP was greater than zero in either the *neuFace_noCFS* or *neuFace* conditions. We used one-sample *t*-tests rather than paired-sample *t*-tests or repeated-measures ANOVAs because we were interested in whether either condition evoked a significant response, not whether the magnitude of any such response varied as a function of condition. For example, a paired-samples *t*-test might show that the SSVEP during conscious perception was larger than during nonconscious perception, but this would not tell us whether the latter evoked a response greater than zero.

Complimentary frequentist one-sample *t*-tests were also performed.

## 3. Results

### 3.1 Study 1: Behavioral results

Across all participants (N=25) the breakthrough time during continuous flash suppression was significantly faster than the full run duration (M_diff_ = 12.59 s, SD = 12.06; Figure 2). A one-sample one-sided *t*-test found that this was a significant difference (*t*(24) = 5.22, *p* < .001, *d* = 1.04). This was true even when we restricted the analysis to include only the subset of participants who were included in the SSVEP analysis (M_diff_ = 8.55 s, SD = 9.21); *t*(18) = 4.05, *p* < .001, *d* = .93).

**Figure 2.**
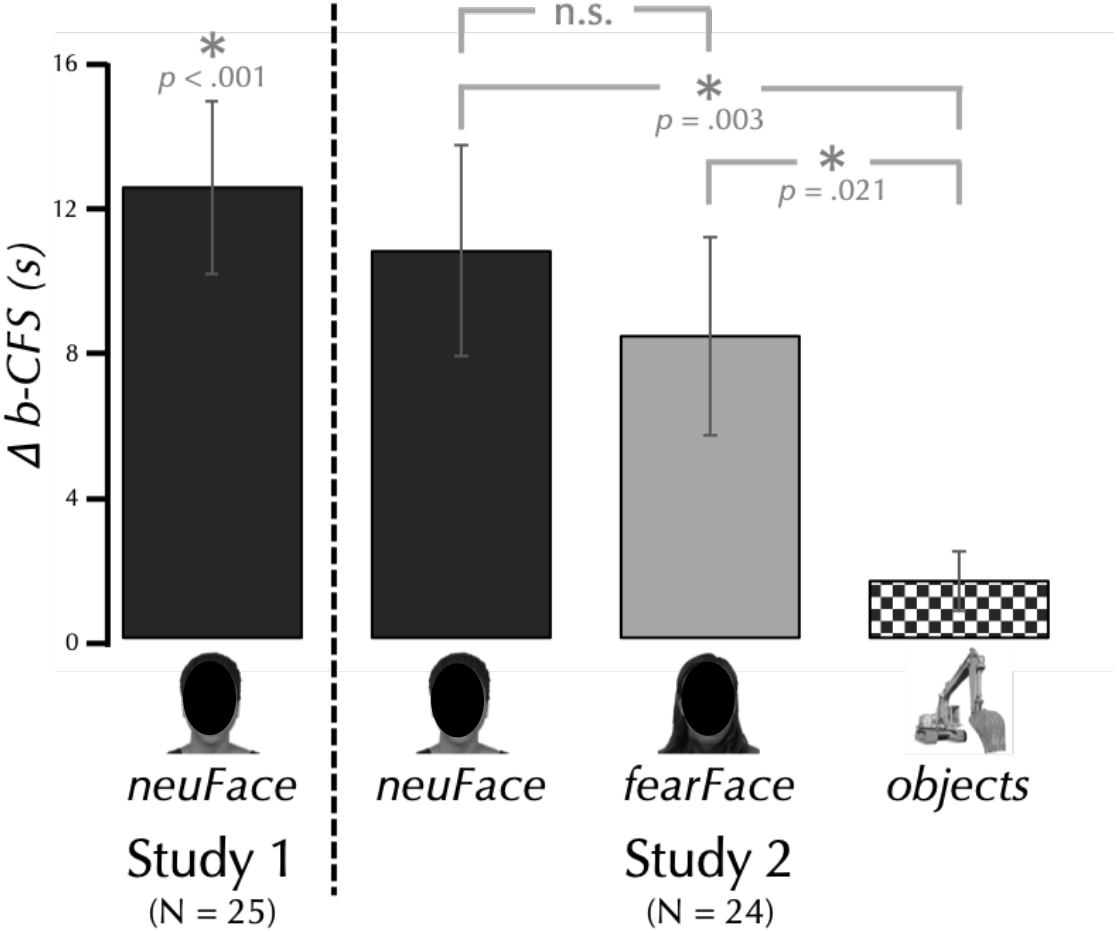
CFS breakthrough time (b-CFS) advantage in Study 1 & 2. (*Face images are obscured for this preprint*). The bar graph shows the difference between the entire possible run duration and the actual average run duration, so a larger number indicates a faster breakthrough time. Note: the b-CFS results from both Study 1 and Study 2 are presented here, but each included an independent sample and was subject to a different analysis. The results of Study 1 (to the left of the vertical dashed line) were analyzed using a one-way one-sample *t*-test. The results of Study 2 show the results of Bonferroni corrected post-hoc tests (see Methods).

### 3.2. Study 1: EEG results

Figure 3 shows the scalp distribution of power at the first and second harmonic, and the average SSVEP to the *CFS* and *noCFS* conditions. The null hypothesis for each condition states that the SSVEP is equal to zero, H_0_: δ = 0. The alternative hypothesis states that effects are positive values and thus δ was assigned a Cauchy prior distribution with r = 1 / √2, truncated to allow only positive effect sizes. For the *neuFace-noCFS* condition (M = .328 μV, SD = .24) we found extreme evidence (BF = 3736) that the observed data are more likely under H_1_ (δ > 0) than under H_0_ (δ = 0). In contrast, for the *neuFace* condition (M = .002 μV, SD = .05) we found moderate evidence (BF = .281) that the results are more likely (specifically, 3.56 times more likely) under H_0_ (δ = 0) than under H_1_ (δ > 0).

**Figure 3.**
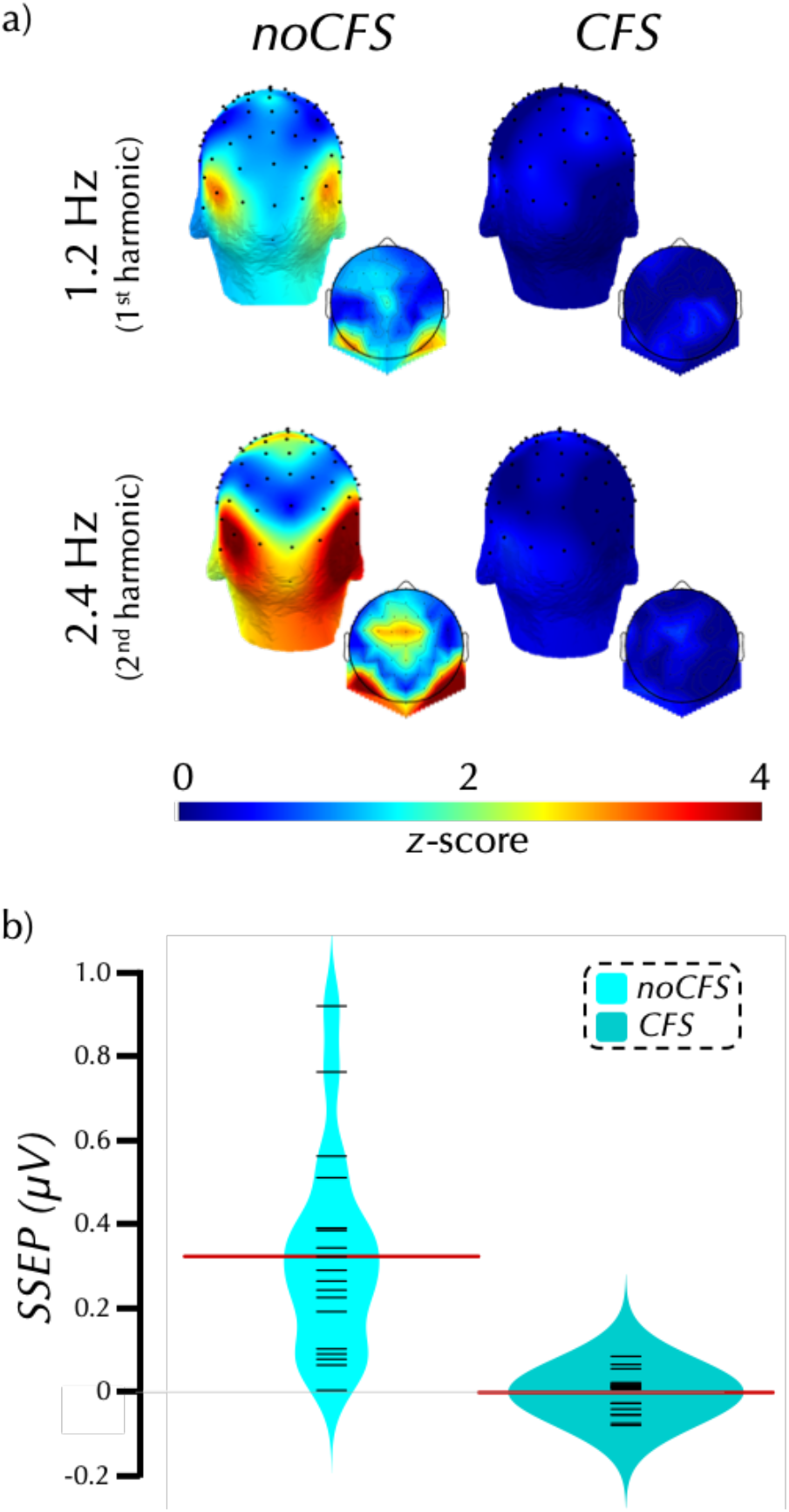
Study 1 SSVEP to neutral faces with and without CFS: These 3-D (larger) and 2-D (smaller inset) scalp maps display the distribution of normalized power at the first (aka fundamental) and second harmonics of the oddball presentation frequency (a). During *noCFS* there was no interocular suppression and the participants were therefore consciously aware of all presented stimuli. During *CFS* there was interocular suppression and the participants were therefore unaware of the stimuli of interest presented to the ‘suppressed’ eye. The bean plots (b) display the average amplitude of the response (μV) combined across the first and second harmonics and across three electrodes of interest: P8, P10, and PO8 for each participant. For each condition, the plot displays the individual participant results (black lines), the distribution density of the results (mirrored across the vertical axis), and the mean response (red line).

The results of the frequentist one-paired *t*-tests were qualitatively the same as the Bayesian tests. A significant response was evoked by *neuFace-noCFS* (*t*(18) = 5.97, *p* < .001, *d* = 1.37), but not *neuFace-CFS* condition (*t*(18) = .22, *p* = .42).

### 3.3. Study 2: Behavioral results

Across all participants (N=24), a one-way repeated-measures ANOVA showed that the breakthrough times significantly varied as a function of condition (*F*(1.67, 38.3) = 9.81, *p* < .001; Figure 2). Bonferroni corrected post-hoc tests showed that *object* breakthrough time (M = 1.58 s, SD 4.1) was significantly slower than for *neuFace* (M = 10.83 s, SD = 14.45, *p* = .003, *d* = .76) and *fearFace* (M = 8.44 s, SD = 13.52, *p* = .021, *d* = .61). Breakthrough times did not did not differ between *neuFace* and *fearFace* (*p* = .463).

A second repeated-measures ANOVA was run on the subset of participants who were included in the SSVEP analysis. However, only 17 of the 22 participants contributed data to all conditions, and therefore the remaining five were held out of this analysis. As with the full sample, breakthrough times significantly varied as a function of condition (*F*(1.97, 31.57) = 4.50, *p* = .019). Bonferroni corrected post-hoc tests showed that *object* breakthrough time (M = 0.18 s, SD 0.75) was significantly slower than for *neuFace* (M = 6.96 s, SD = 10.24, *p* = .038, *d* = .68), but not *fearFace* (M = 4.51 s, SD = 9.82, *p* = .221). Breakthrough times did not differ between *neuFace* and *fearFace (p* = .838).

### 3.4. Study 2: EEG results

Figure 4 shows the scalp distribution of power at the first and second harmonic, and the average SSVEP to the *CFS* and *noCFS* conditions. The null hypothesis for each of the Bayesian one-sample *t*-tests were as described for Study 1. For all three *noCFS* we found extreme evidence the observed data are more likely under H_1_ (δ > 0) than under H_0_ (δ = 0): *neuFace_noCFS* (M = .353 μV, SD = .23, BF = 30,161), *fearFace_noCFS* (M = .305 μV, SD = .17, BF = 72,251), *object_noCFS* (M = .573 μV, SD = .31, BF = 290,318). In contrast, for each of the *CFS* conditions we found anecdotal to moderate evidence that the observed data are more likely under H_0_ (δ = 0) than under H_1_ (δ > 0): *neuFace_noCFS* (M = −.019 μV, SD = .17, BF = .167), *fearFace_noCFS* (M = .022 μV, SD = .10, BF = .614), *object_noCFS* (M < .001 μV, SD = .06, BF = .243). Figure 3 shows the scalp distribution of power at the first and second harmonic, and the average SSVEP to the *CFS* and *noCFS* conditions.

**Figure 4.**
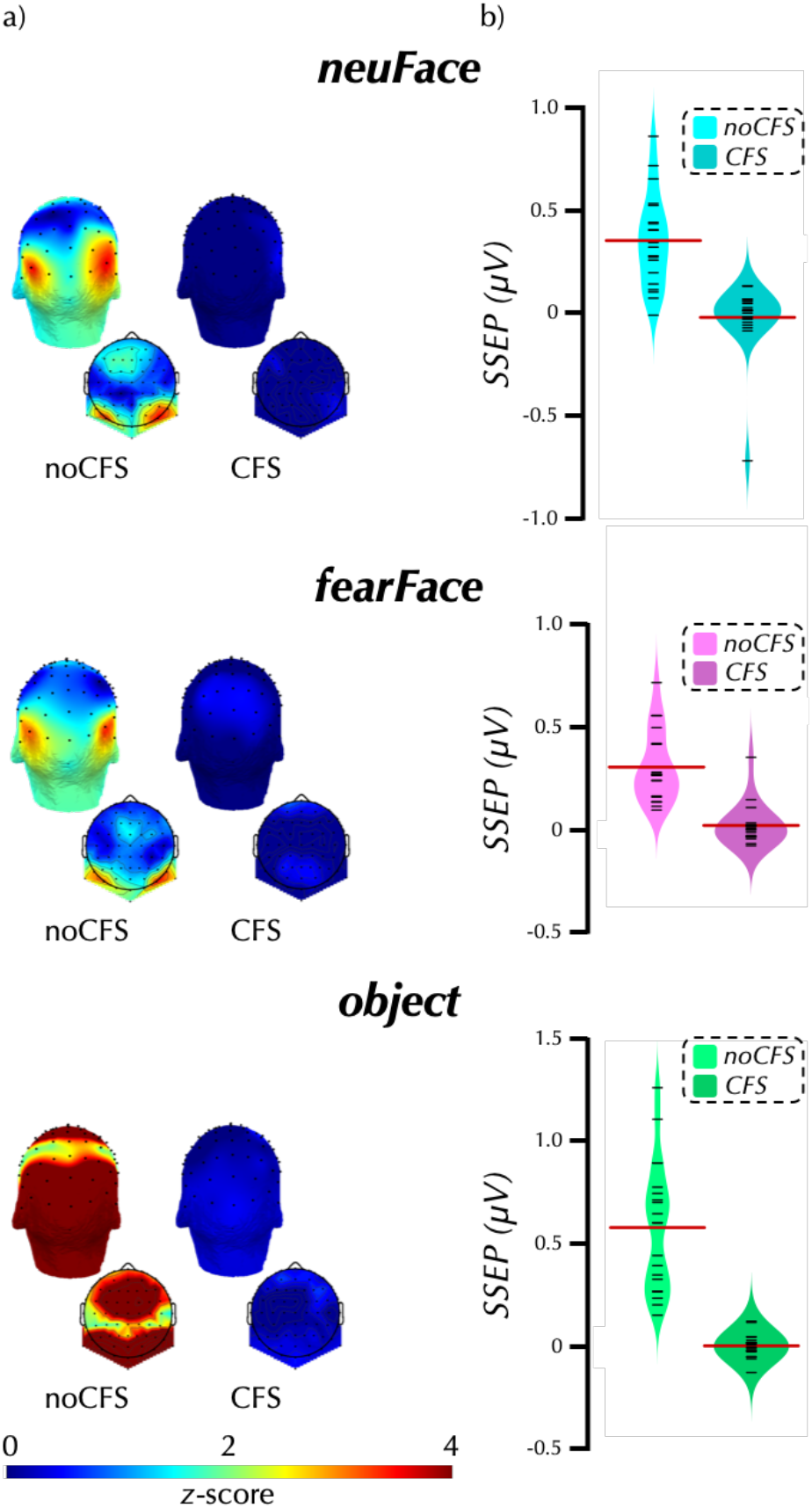
Study 2 SSVEP: These 3-D (larger) and 2-D (smaller inset) scalp maps display the distribution of normalized power at the first harmonic (aka fundamental) of the oddball presentation frequency (a). During *noCFS* there was no interocular suppression and the participants were therefore consciously aware of all presented stimuli. During *CFS* there was interocular suppression and the participants were therefore unaware of the stimuli of interest presented to the ‘suppressed’ eye. The bean plots (b) display the average amplitude of the response (μV) combined across the first and second harmonics and across three electrodes of interest: P8, P10, and PO8 for each participant. For each condition, the plot displays the individual participant results (black lines), the distribution density of the results (mirrored across the vertical axis), and the mean response (red line).

The results of the frequentist one-sample *t*-tests were qualitatively the same as the Bayesian tests. A significant response was evoked by all of the *noCFS* conditions: *neuFace-noCFS* (*t*(19) = 6.97, *p* < .001, *d* = 1.56), *fearFace-noCFS* (*t*(19) = 7.65, *p* < .001, *d* = 1.75), *object-noCFS* (*t*(19) = 8.27, *p* < .001, *d* = 1.85). In contrast, the *CFS* conditions did not yield any significant effects (*ps* ≥ .33).

### 3.5. Combined Study 1 and Study 2 EEG results

In order to maximize SNR and thus detection sensitivity, we analyzed the combined *neuFace* SSVEP data from Study 1 and Study 2 resulting in a larger sample of N = 39. For the *neuFace_noCFS* condition (M = .341 μV, SD = .23) we found extreme evidence (BF = 6.699e+8) that the observed data are more likely under H_1_ (δ > 0) than under H_0_ (δ = 0). In contrast, for the *neuFace* condition (M = −.009 μV, SD = .13) we found moderate evidence (BF = .128) that that observed the results are more likely (specifically, 7.81 times more likely) under H_0_ (δ = 0) than under H_1_ (δ > 0).

The results of the frequentist one-sample *t*-tests were qualitatively the same as the Bayesian tests. A significant response was evoked by *neuFace-noCFS* (*t*(38) = 9.25, *p* < .001, *d* = 1.48), but not *neuFace-CFS* condition (*t*(38) = .43, *p* = .67).

### 3.6. Study3: Behavioral results

Across all participants (N=35), a one-way repeated-measures ANOVA showed that breakthrough time was significantly affected by condition (*F*(2.81, 95.57) = 3.60, *p* = .018; Figure 5). One-way paired-samples *t*-tests were used for four planned comparisons. The breakthrough time for *faces* (M = 15.56 s, SD 14.52) was significantly faster than for *objects* (M = 10.60 s, 14.53; *t*(34) = 1.71, *p* = .048, *d* = .29), but not *curvilinear* objects (M = 18.69 s, SD 15.57; *p* = .910). The breakthrough time for *curvilinear* objects was significantly faster than for both *objects* (*t*(34) = 2.81, *p* = .004, *d* = .48) and *rectilinear* objects (M = 11.87 s, 12.99; *t*(34) = 2.39, *p* = .011, *d* = .40).

**Figure 5.**
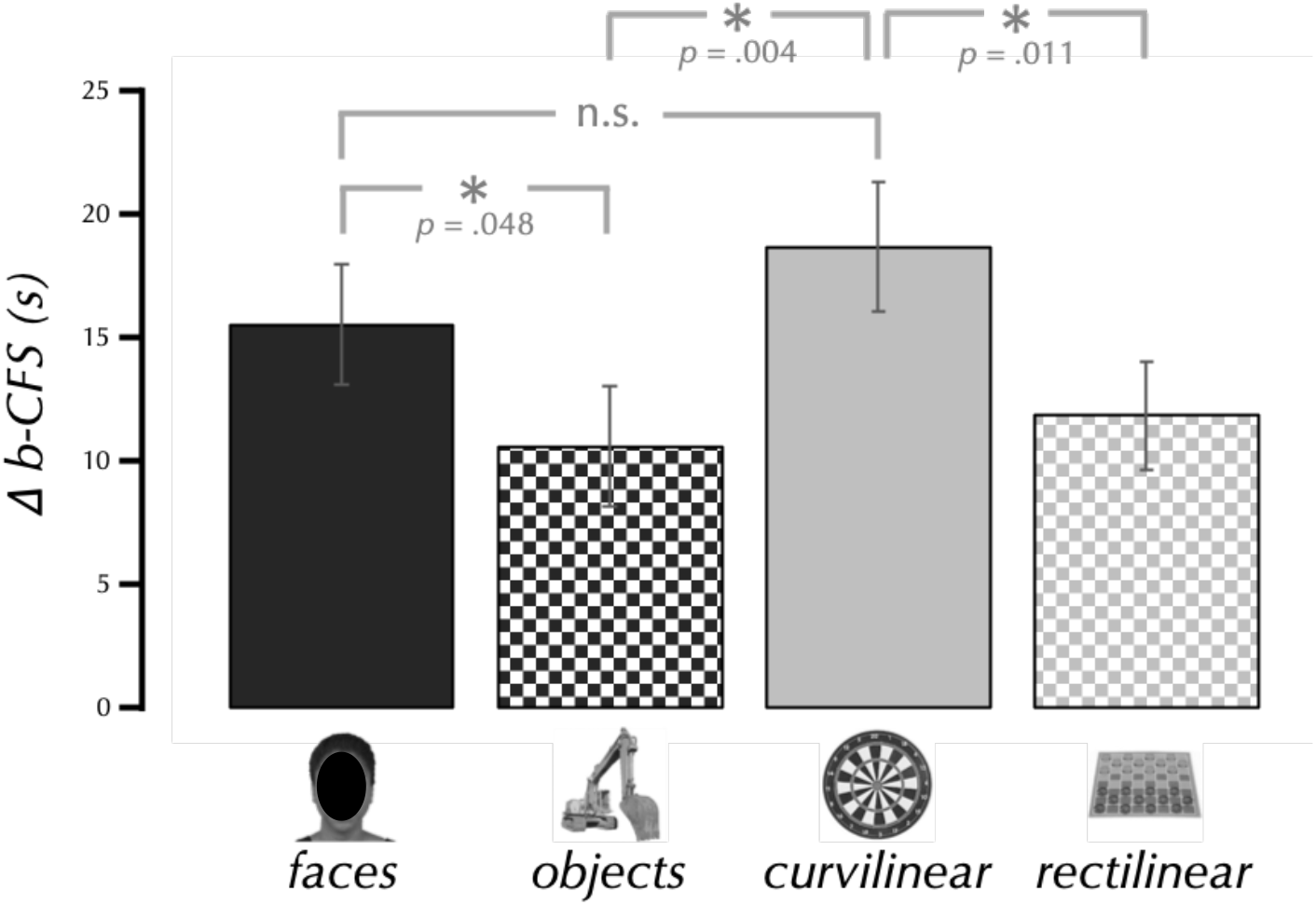
Breakthrough time advantage in Study 3. (*Face images are obscured for this preprint*). The bar graph shows the difference between the entire possible run duration and the actual average run duration. Thus, a larger number indicates a faster b-CFS. The data (N = 35) were analyzed with four planned one-way paired-samples *t*-tests (see Methods).

## 4. Discussion

We report two experiments that do not support the engagement of cortical face-selective regions during interocular suppression. A face-sensitive steady-state visually evoked potential (SSVEP) to neutral faces (Studies 1 & 2) or fearful faces (Study 2) was observed only when participants were aware of the stimuli. In contrast, we observed evidence of selective nonconscious processing; faces broke through interocular suppression faster than objects. We followed up on these results with a third study in which we observed faster breakthrough time for curvilinear than rectilinear objects. Moreover, the breakthrough time for curvilinear objects did not differ from neutral or fearful faces. We interpret these results as follows: 1) evidence that cortical face-selective regions are not engaged during face perception without awareness, 2) this is true for fearful, as well as neutral, faces, and 3) faster breakthrough times for faces is owed to the curvilinearity common to all faces rather than to high-level category membership.

### 4.1 No EEG evidence of nonconscious detection of neutral faces

In Studies 1 and 2 we did not find an EEG response indicating face detection when faces were presented outside of conscious awareness. At first blush, this finding is simply another result added to a conflicted literature (for a recent review, see Axelrod et al., 2015) in which some have found M/EEG signals associated with nonconscious detection (Henson, Mouchlianitis, Matthews, & Kouider, 2008; Sterzer et al., 2009; Jiang et al., 2009; Suzuki & Noguchi, 2013), whereas others have not (Reiss & Hoffman, 2007; Harris et al., 2011; Navajas et al., 2013; Shafto & Pitts, 2015; Kume et al., 2016). But, our unique combination of CFS and SSVEP addresses at least three limitations of the prior studies and therefore represent a meaningful contribution to the literature.

First, it is possible that the negative reports simply failed to detect a noisy, but nonetheless present, face-selective response. Here, we address that issue by taking advantage of the high signal-to-noise ratio of SSVEP (Norcia et al., 2015). Despite the higher SNR afforded by SSVEP, our results are consistent with those prior reports that failed to find an EEG marker of face processing without awareness. Moreover, we used a Bayesian analysis to test the probability of the null model given the data, rather than simply testing the probability of observing an effect if the null were true. This support for the null model is easily understood with a simple visual inspection of the scalp distributions of power at the face presentation frequency (Figures 3 and 4), which show no hint of an SSVEP.

Second the inconsistent findings may be owed to the different blinding methods used across experiments. Axelrod, Bar, and Rees (2015) found that EEG studies that report no evidence of nonconscious face processing tend to use variations of masking paradigms, whereas those that find evidence tend to use variations of dichotic stimulation (but see Izatt, Dubois, Faivre, & Koch, 2014; Shafto & Pitts, 2015). This might suggest a partial awareness during CFS (Mudrik, Gelbard-Sagiv, Faivre, & Koch, 2013; Gayet, Van der Stigchel, & Paffen, 2014; Stein & Sterzer, 2014) that results in false positives. Here, we report evidence against face detection without awareness, despite using an approach that is ostensibly more likely to produce a positive result.

Third, the inconsistent findings may be owed to variation in what is considered “awareness” across studies (Faivre, Berthet, & Kouider, 2014; Peters & Lau, 2015). As with the blinding differences noted above, the concern is that differences in instructions and/or participant response biases (Rodríguez et al., 2012) could lead to false positives. That is, if participants employ a conservative detection criterion, they might view images with some degree of conscious awareness without reporting it. The current work is less susceptible to such bias because SSVEP requires continuous periodic stimulation. Therefore, a false positive would require that participants experience unreported awareness for extended durations rather than just on discrete events. In the current work, general response bias is even less likely given that participants detected faces faster than objects (Studies 2 and 3) and curvilinear objects faster than rectilinear objects (Study 3). Perhaps most importantly, any concern that a response bias might lead to Type I errors should be assuaged by the fact that we did not observe a positive SSVEP during periods which the participants did not report awareness.

### 4.2. No evidence of nonconscious detection of fearful faces

Are emotionally relevant signals privileged relative to neutral signals? It has been proposed that affective signals are qualitatively different than neutral signals and processed via subcortical pathways (Tamietto & de Gelder, 2010; but see Pessoa & Adolphs, 2010) or, in the case of face processing, cortical pathways distinct from those that support identity processing (Haxby, Hoffman, & Gobbini, 2000; but see Calder & Young, 2005). Faivre, Berthet, and Kouider (2014) note that there is more consistent evidence for nonconscious processing of facial expression than there is for facial identity, and therefore that “The discrepancy between the processing of facial identity and facial expressions suggests that the latter may be processed along subcortical routes that are not fully disrupted by CFS” (p. 8).

Support for an affective advantage comes primarily from fMRI and behavioral studies (for reviews see Sterzer, Stein, Ludwig, Rothkirch, & Hesselmann, 2014; Axelrod et al., 2015; Diano, Celeghin, Bagnis, & Tamietto, 2016), though there are also a handful of EEG reports (Jiang et al., 2009; Jessen & Grossmann, 2014). However, some have argued that carefully controlling for potential methodological confounds causes any affective advantage to disappear (Straube, Dietrich, Mothes-Lasch, Mentzel, & Miltner, 2010; Pessoa & Adolphs, 2010; Hoffmann, Mothes-Lasch, Miltner, & Straube, 2015; Hedger, Adams, & Garner, 2015; Hedger, Gray, Garner, & Adams, 2016). Schlossmacher, Junghöfer, Straube, and Bruchmann (2017) found that modulation of face-sensitive ERPs (e.g., N170) observed during conscious perception were not observed during interocular rivalry with CFS. Finally, the authors of a recent metaanalysis of behavioral experiments conclude “uncritical acceptance of the standard hypothesis, which states that threat stimuli can be identified and prioritized without awareness, is premature.” (Hedger et al., 2016, p. 961). But notably, they report that fearful faces were the only threat stimulus that consistently showed evidence of a nonconscious advantage in b-CFS paradigms. Therefore, we must entertain the possibility that there is an affective advantage in nonconscious processing with a source that is not readily detectable with EEG.

### 4.3. Is there a subcortical effect?

A limitation of EEG, and thus the current work, is that potentials generated in subcortical structures will have lower SNR due to their increased distance from recording sites on the scalp. For some structures, such as the amygdala, this is exacerbated by a spatial organization of neurons that results in local volume currents cancelling each other out rather than summating into a field large enough to be detected on the scalp (Silva, 2018). The current pattern of results – faster breakthrough times despite the absence an of EEG signature – would be consistent with faces being processed nonconsciously by subcortical systems and would explain the behavioral advantage without concomitant SSVEP. This was our initial conclusion after seeing the results of Study 1. But the full pattern of results across all three studies makes this unlikely for at least two reasons.

First, if a subcortical pathway existed for fast processing and thus attentional orienting, one would reasonably assume that this system would engage the relevant cortical systems that are specialized for processing the to-be-attended stimuli (Brooks et al., 2012). In contrast, we found no indication of cortical engagement. Though we note that there are studies which have found evidence of amygdala activation without concomitant cortical activation (see de Gelder, van Honk, & Tamietto, 2011). Second, and perhaps more important, we did not observe a faster breakthrough time for fearful faces compared to neutral faces. If anything, the breakthrough time for fearful faces was slower (though this was a small and insignificant difference). So, on the one hand, the current data cannot rule out subcortical nonconscious processing of fearful faces. On the other hand, if such processing occurs, it does so without engaging cortical face processing systems and without conferring an observable behavioral advantage.

### 4.4. The effect of mid-level visual features on b-CFS

At first blush, the EEG and behavioral results of Studies 1 and 2 seem incompatible. We observed a significantly faster breakthrough time for faces than objects, but no face-selective SSVEP. We believe the results of Study 3 – faster breakthrough times for curvilinear than for rectilinear objects – elucidate the nature of this contradiction. Specifically, we interpret these results as evidence that the behavioral advantage for face processing is owed to the curvilinearity of faces rather than their high-level category membership. This interpretation is consistent with a growing literature that focuses on the importance of mid-level feature processing in the visual system.

Perhaps most relevant to the current work is a recent study by Moors, Wagemans, and de-Wit (2016) in which b-CFS was investigated as a function of curvature relative to fixation. Participants viewed the left half or the right half of a face in either an upright or inverted orientation presented to the left or right of fixation. Thus, faces were either presented with natural convex, or unnatural concave curvature relative to fixation. They found that curvature relative to fixation played an important role in faster breakthrough such that natural convexity was faster than concavity. This is consistent with prior work that found a preference for convex contours in area V4 of the macaque (Pasupathy & Connor, 1999). In the current studies, all curvilinear images (faces and objects) were convex relative to fixation, so the data cannot speak to the importance of convexity vs. concavity, but do support priority for curvilinear over rectilinear contours.

We presented all stimuli roughly centered at fixation, thus resulting in processing occurring primarily in regions of visual cortex with foveal and parafoveal receptive fields. This might contribute to the observed nonconscious preference for curvilinear shapes. In macaque visual cortex there is a correlation between contour and eccentricity such that curvilinear contours are preferred in the central visual field, whereas rectilinear contours are preferred in the periphery (Srihasam, Vincent, & Livingstone, 2014). This observed relationship is particularly strong in early visual cortex, but a general preference for curvature has been observed to increase from lower to higher visual processing areas (Wilkinson et al., 2000; Ponce, Hartmann, & Livingstone, 2017) and might contribute to the organization of high-level visual cortex (Nasr, Echavarria, & Tootell, 2014; Srihasam et al., 2014; Andrews, Watson, Rice, & Hartley, 2015; Long, Yu, & Konkle, 2018).

Human face selective regions are particularly sensitive to curvilinearity (Caldara et al., 2006). Indeed, prosopagnosia (aka face-blindness) seems to selectively impair processing of curved edges and shapes (Kosslyn, Hamilton, & Bernstein, 1995). Similarly, a network of curvature sensitive regions in the macaque brain is adjacent to face-sensitive regions suggesting a possible functional relationship (Yue, Pourladian, Tootell, & Ungerleider, 2014). There is also an intriguing relationship between curvilinearity and animacy such that behavioral categorization is largely dependent on the amount of curvilinearity present in the image with images of animate things being more curvilinear than images of inanimate things (Long, Störmer, & Alvarez, 2017; Zachariou, Del Giacco, Ungerleider, & Yue, 2018; but also see Proklova, Kaiser, & Peelen, 2016). Furthermore, a recent ERP study found evidence that animals and non-animals were distinguished nonconsciously (Zhu, Drewes, Peatfield, & Melcher, 2016).

Do mid-level features, particularly curvilinearity, account for observed differences in b-CFS paradigms? Our results are consistent with prior reports that would suggest so. Such features have been shown to drive the nonconscious processing of face identity (Gelbard-Sagiv, Faivre, Mudrik, & Koch, 2016), expression (Hedger et al., 2015), and dominance (Stein, Awad, Gayet, & Peelen, 2018; Gayet et al., 2014). Though it is yet unclear whether these features are themselves being processed nonconsciously (Pitts, Martínez, & Hillyard, 2012), or if the effect is due to partial awareness (Gelbard-Sagiv et al., 2016).

### 4.5. Two-threshold model

The contribution of mid-level features discussed above might account for many of the studies that have reported nonconscious processing of several different dimensions of face perception (for review see Axelrod et al., 2015), but others are less easily explained. For example, Gobbini and colleagues (2013) report that faces oriented directly toward the viewer breakthrough faster than faces oriented slightly away. In this case, both conditions have near-identical curvilinearity and, importantly, convexity relative to fixation. What might drive this effect if not mid-level features?

One intriguing possibility is that nonconscious processing is not an all-or-none phenomenon, but rather can be considered a process of degree. This “two-threshold model of nonconscious processing” (Schlossmacher et al., 2017) posits that some features might only be processed when in the shallow depths of unconsciousness (Peremen & Lamy, 2014; Sterzer et al., 2014). In the context of this model, mid-level features might push faces from the depths toward the waterline of consciousness, at which point they are susceptible to privileged processing that ultimately causes a faster breakthrough. On the one hand, our results can be interpreted as being broadly consistent with such a model. On the other hand, we did not observe an advantage of fearful faces compared to neutral faces, or neutral faces compared to curvilinear objects. In other words, we did not observe an additive benefit of high-level category membership (face vs. object) beyond what could be explained by mid-level features (curvilinear vs. rectilinear).

### 4.6. Limitations

We have addressed several limitations of the current work in the prior discussion. Here we will briefly address three more. First, we used low opacity images (see Methods) to extend suppression time. It is possible this accounts for our inability to detect an EEG response, but this seems unlikely because we did observe a behavioral effect despite the low opacity. It should also be noted that the low opacity images evoked a sufficient signal in the *noCFS* conditions. Second, faces are a substantially more homogenous set of stimuli than are objects. It is possible that the repetition of homogenous oddballs facilitated a faster breakthrough time. We think this is unlikely given the design and the results of Study 3. In that study, we observed a faster breakthrough time for curvilinear objects than for rectilinear objects, despite there being no appreciable difference in the homogeneity of the seven exemplars within each condition. Third, it is possible that the observed curvilinearity effect was owed to contrast with the rectilinear continuous flash stimuli. We cannot exclude this possibility because these studies do not include a version in which the CFS stimuli are curvilinear. The substantial evidence for the importance of curvilinearity in both low-level and category-selective regions of the visual system leads us to conclude that this explanation is unlikely. Further, both faces and curvilinear objects were suppressed with rectilinear CFS stimuli, so this could not account for the important observation that the breakthrough times did not differ for faces and curvilinear objects.

### 4.7. Conclusions

The results of these studies suggest that cortical face-selective regions do not engage in nonconscious face processing. Moreover, the observed advantage faces have over non-faces in breaking through flash suppression is likely due to their curvilinearity, rather than their high-level category membership. In the current series of studies, we were unable to find EEG evidence in support of the notion that faces are processed without benefit of conscious awareness. Paradoxically, we did observe a faster breakthrough time into conscious awareness for faces than for objects. Faster b-CFS is commonly interpreted to indicate nonconscious processing (Jiang et al., 2007). In a follow-up study, we found evidence that the mid-level visual features of a face – specifically, their curvilinearity – account for the faster breakthrough time, rather than their high-level category membership, and therefore are why we see no EEG signature of face perception.

## 5. Acknowledgments

We thank Sabrina Halavi and Josue Parr for their contributions to this work. We also thank Sarah Mohr and Anxu Wang for help in data collection. This research was funded, in part, by the Kenyon Summer Science Program.

